## Supplemental data for "Zebrafish Polymerase Theta and human Polymerase Theta: orthologues with homologous function"

Supplemental data 1

CLUSTAL O(1.2.4) zPol θ and hPol θ alignment

Red indicates loop insertions

```
hPolQ      GFKDNSPISDTSFSLQLSQDGLQLTPASSSSESLSIIDVASDQNLFQTFIKEWRCKKRFS	60
zPolQ      -----------------------------------IIDVASDRRLFETFVNEWKTKERFS	25
                                              *******:.**:**::**: *:***

hPolQ      ISLACEKIRSLTSSKTATIGSRFKQASSPQEIPIRDDGFPIKGCDDTLVVGLAVCWGGRD	120
zPolQ      LAVACEKTDSTSVQPETVIGGKFKKPTTPMR-NKRKDGFLLKGYEDLVVIGISVSWGAKD	84
           :::****  * : .  :.**.:**: ::* .   *.*** :** :* :*:*::*.**.:*

hPolQ      AYYFSLQKEQKHSEISASLVPPSLDPSLTLKDRMWYLQSCLRKESDKECSVVIYDFIQSY	180
zPolQ      AYFVSLQQELVDTDISASLAPPPLDDTLTVEERLKQIQSCLQKDSS---VTVTYDFIHLY	141
           **:.***:*  .::*****.** ** :**:::*:  :****:*:*.    .* ****: *

hPolQ      KILLLSCGISLEQSYEDPKVACWLLDPDSQEPTLHSIVTSFLPHELPLLEGMETSQGIQS	240
zPolQ      KILLLACELAVRGTFEDPKIACWLLDSSSKERTLHNMVTSFATEDLPMLEGISAGQGVQS	201
           *****:* :::. ::****:****** .*:* ***.:****  .:**:***:.:.**:**

hPolQ      LGLNAGSEHSGRYRASVESILIFNSMNQLNSLLQKENLQDVFRKVEMPSQYCLALLELNG	300
zPolQ      LGIYGEASQPGRYRAAIESVLVFRVMTQLNCLLEKDGFLDVFKKVEMPTQYCLALLELNG	261
           **: . :.: *****::**:*:*. *.***.**:*:.: ***:*****:***********

hPolQ      IGFSTAECESQKHIMQAKLDAIETQAYQLAGHSFSFTSSDDIAEVLFLELKLPPNREMKN	360
zPolQ      IGFSIAECEAQKHVMQAKLSALESQAYQLAGHSFSLTSPEDVAEVLFLELKLPPNGDLNG	321
           **** ****:***:*****.*:*:***********:** :*:************* :::.

hPolQ      QGSKKTLGSTRRGIDNGRKLRLGRQFSTSKDVLNKLKALHPLPGLILEWRRITNAITKVV	420
zPolQ      LKNKKTLGYTRR---AGARIKLSKQFSTTKDVLEKLKPLHPLPGVILEWRRITNALTKVV	378
             .***** ***    * :::*.:****:****:*** ******:**********:****

hPolQ      FPLQREKCLNPFLGMERIYPVSQSHTATGRITFTEPNIQNVPRDFEIKMPTLVGESPPSQ	480
zPolQ      FPLQREKKWHSHLKMDRIHPISQSHTATGRVSFTEPNIQNVPKDFEIQMPTLIEESQTSQ	438
           *******  : .* *:**:*:*********::**********:****:****: **  **

hPolQ      AVGKGLLPMGRGKYKKGFSVNPRCQAQMEERAADRGMPFSISMRHAFVPFPGGSILAADY	540
zPolQ      NGGSKMWCKR-TKINR--LL--APLLKVSDKSPDKGMQFSVSMRHAFVPFSGGLILAVDY	493
             *. :      * ::   :      ::.::: *:** **:********* ** ***.**

hPolQ      SQLELRILAHLSHDRRLIQVLNTGADVFRSIAAEWKMIEPESVGDDLRQQAKQICYGIIY	600
zPolQ      SQLELRILAHLSRDRRLLHVLNSGADVFKSIAAEWKMVDPASVDDNMRQQAKQICYGIIY	553
           ************:****::***:*****:********::* **.*::*************

hPolQ      GMGAKSLGEQMGIKENDAACYIDSFKSRYTGINQFMTETVKNCKRDGFVQTILGRRRYLP	660
zPolQ      GMGAKSLGEQMGIEENDAACYIETFKSRYNGIQNFLRETVQKCGKNGYVKTLLGRKRFLP	613
           *************:********::*****.**::*: ***::* ::*:*:*:***:*:**

hPolQ      GIKDNNPYRKAHAERQAINTIVQGSAADIVKIATVNIQKQLETFHSTFK-SHGHREGMLQ	719
zPolQ      GIKDSNVYIKSHAERQAVNTTVQGSAADIVKLATINIQRRIEEAFPGVPTSHQHP-----	668
           ****.* * *:******:** **********:**:***:::*  .  .  ** *

hPolQ      SDRTGLSRKRKLQGMFCPIRGGFFILQLHDELLYEVAEEDVVQVAQIVKNEMESAVKLSV	779
zPolQ      ----SIRLGGRHRNQFRPLRGGYFILQLHDELIYEVAEEDVIQVAQIVKREMESVVKLYV	724
               .:    : :. * *:***:*********:********:*******.****.*** *

hPolQ      KLKVKVKIGASWGELKDFDV	799
zPolQ      KLRVKVKVGPSWGNLQDLDI	744
           **:****:* ***:*:*:*:
```

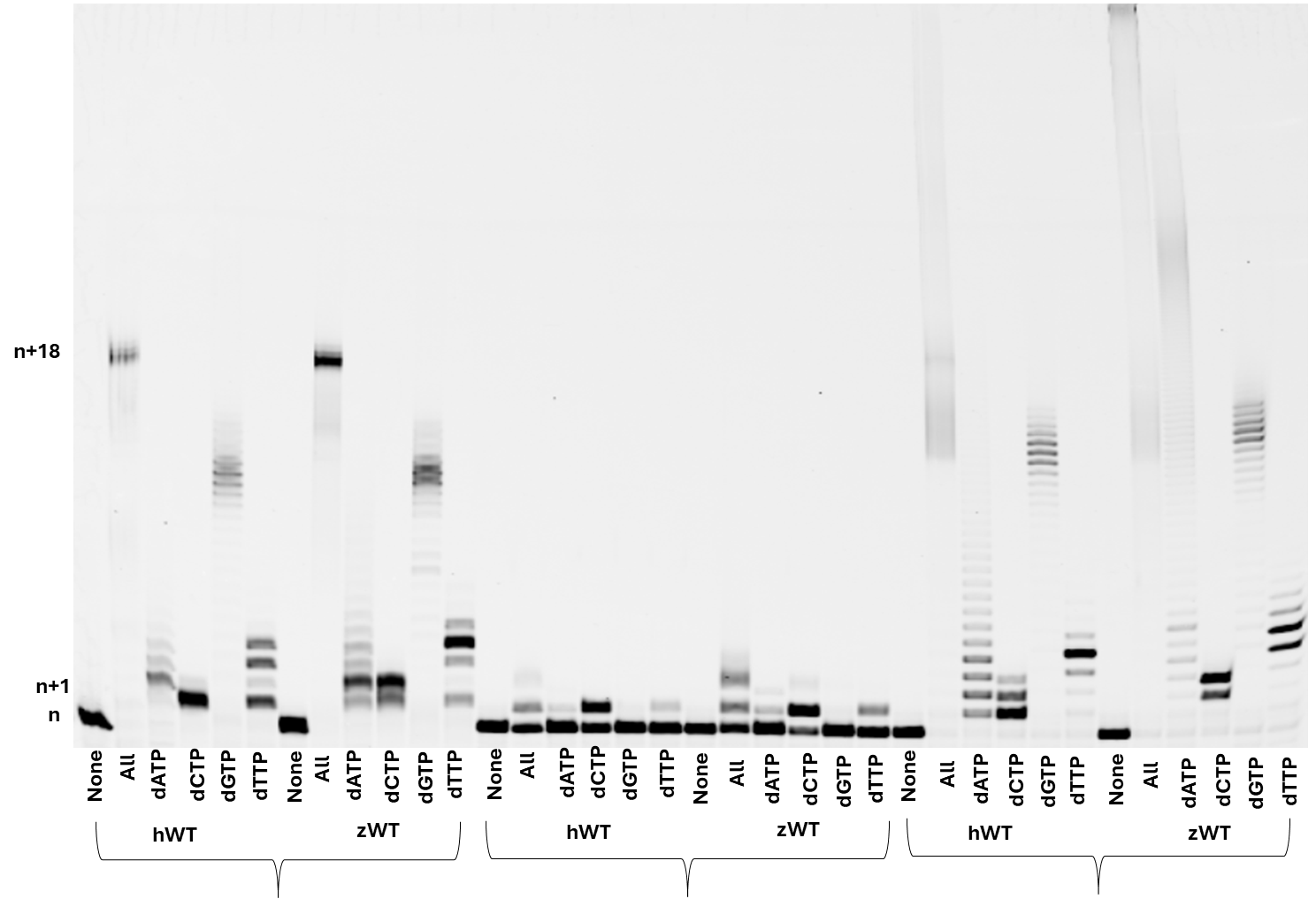

**Mg^2+^**

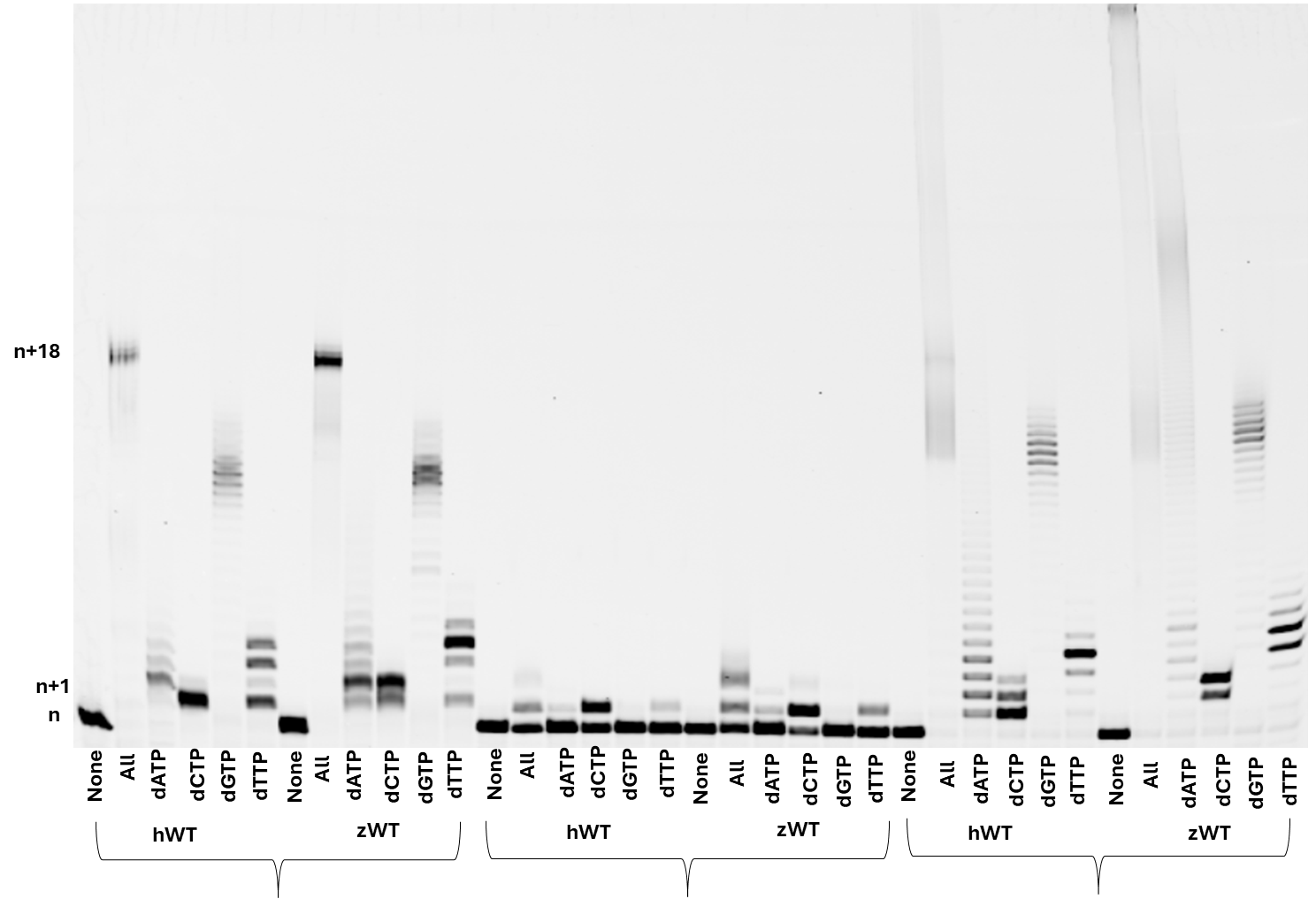

**Mn^2+^**

**Supplement 2.** Both hWT and zWT Pol θ were assayed under single-turnover conditions at t 4:1 ratio protein:DNA (see Materials and Methods). Pol θ and 25/40 dsDNA were preincubated and combined with either 10 mM MgCl_2_ or MnCl_2_ for 5 minutes and 37°C. DNA extension products were separated on a denaturing gel and visualized on a Typhoon scanner.

**Mn^2+^**
